# In vitro determination of inhibitory effects by humic substances complexing Zn and Se on SARS-CoV-2 virus replication

**DOI:** 10.1101/2021.08.11.456012

**Authors:** Polett Hajdrik, Bernadett Pályi, Zoltán Kis, Noémi Kovács, Daniel S. Veres, Krisztián Szigeti, Imre Hegedűs, Tibor Kovács, Ralf Bergmann, Domokos Máthé

**Affiliations:** Department of Biophysics and Radiation Biology, Semmelweis University, Budapest, Hungary; National Biosafety Laboratory, National Public Health Center, Budapest, Hungary; Department of Microbiology, Semmelweis University, Budapest, Hungary; CROmed Translational Research Ltd., Budapest, Hungary; Institute of Radiochemistry and Radioecology, University of Pannonia, Veszprém, Hungary; Helmholtz-Zentrum Dresden Rossendorf, Dresden, Germany; Hungarian Centre of Excellence for Molecular Medicine, In Vivo Imaging Advanced Core Facility, Semmelweis University Site, Szeged, Hungary

**Keywords:** SARS-CoV-2, humic acid, fulvic acid, Zn-Se-ascorbic acid complex, antiviral activity, RT-PCR

## Abstract

Humic substances are well known human nutritional supplement materials and play important performance-enhancing roles as animal feed additives, too. For decades, ingredients of humic substances have also been proven to carry potent antiviral effects against different viruses. Here, the antiviral activity of a humic substance containing ascorbic acid, Se^-^ and Zn^2+^ ions intended as a nutritional supplement material was investigated against SARS-CoV-2 virus B1.1.7 Variant of Concern (“Alpha Variant”) in a VeroE6 cell line. Results show that this combination has a significant *in vitro* antiviral effect at a very low concentration range of its intended active ingredients. Even picomolar concentration ranges of humic substances, vitamin C and Zn/Se ions in the given composition were enough to achieve fifty percent viral replication inhibition in the applied SARS-CoV-2 virus inhibition test.

## 1. Introduction

Humic substances (HS), composed by hymatomelanic acid, humic acids (HA), fulvic acid (FA), ulmic acid and trace minerals are widely known basic components of nutritional supplement materials in human. A large amount of HS are generated in forests and peat. HS have a ubiquitous presence and HS are one of the largest carbon sources in nature. HS originate from decayed plants in the soil that are decomposed by microbes. HS are soluble in alkaline media, partially soluble in water and insoluble in acidic media. The exact composition of HS may differ based on their origin and type of extraction technology. One of their most important components, FA is water-soluble thanks to its polyphenolic molecular structure (Figure 1). Water molecules stabilize its electronic structure even in neutral form (Alvarez-Puebla *et al*., 2006). Fulvic acid (FA) forms small aggregates at pH 5 on gold surfaces and presumably in soil microstructures too. These aggregates compose a fractal-like structure at pH 8 caused by electrostatic forces according to surface-enhanced Raman spectroscopy measurements (SERS) (Alvarez-Puebla *et al*., 2004). The average molar mass of FA is highly dependent of its origin. Native FA is usually a mixture of different molecules (Lieke *et al*., 2021). The molecular weight of native FA (e.g. originating from Suwanee River) is about 2310 Da as provided by literature and the International Humic Substances Society (IHSS) (Chon *et al*., 2013; Chin *et al*., 1994). The other important and well studied component, HA is larger and more complex in structure with much less solubility. The molecular weight of HA changes from 3160 to 26400 Da depending on the origin of the samples (Asakawa *et al*., 2008). Contrary to the molecular weight differences, no significant difference is detectable between the structural building block elements of FA and HA (Schellekens *et al*., 2017; Stevenson, 1994; Sutton and Sposito, 2005; Nebbioso and Piccolo, 2013).

**Figure 1.**
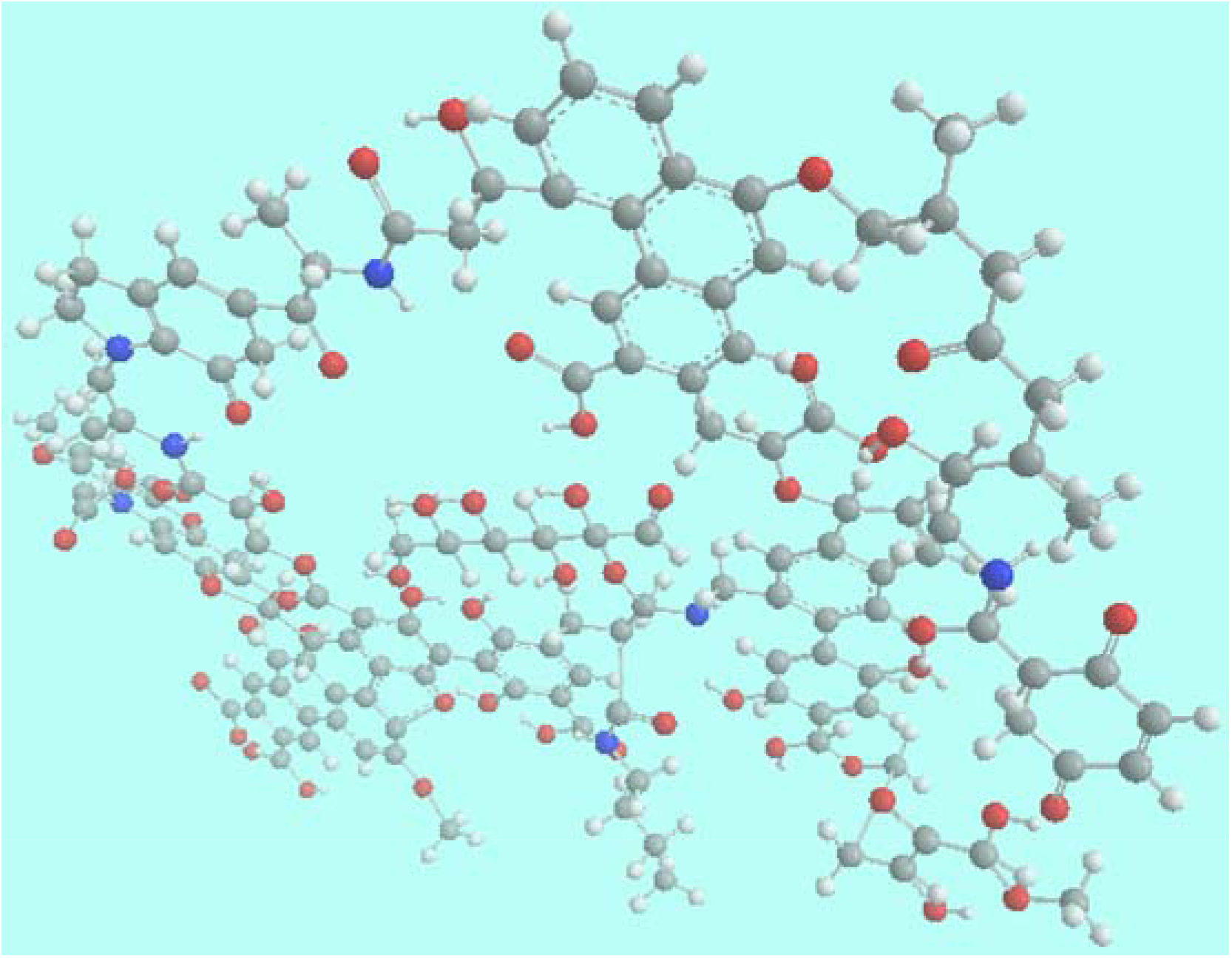
3D molecular structure of a small, characteristic part of HS. This graphícs contains typical structural elements and bonds of natural HS that build their molecular structure randomly based on NMR and IR studies (Lag *et al*., 2008). Grey, carbon; white, hydrogen; blue, nitrogen; red, oxygen atoms. (Graphics was created by ChemDraw Ultra and Chem3D software.)

HS have been used for thousands of years in human healthcare. Humic substances are well-known drugs in Indian Ayurvedic medicine as “shilajit” (Ghosal *et al*., 1991; Ghosal *et al*., 1995; Pant *et al*., 2012; Mishra *et al*., 2019) since time immemorial, as well as in European mud- and balneotherapy from ancient times (Kumar Gautam *et al*., 2021). Shilajit contains mainly FA, HA and trace minerals. Numerous studies show that it has anti-inflammatory, antioxidant, antimutagenic, antiviral, heavy metal chelating, antitumor, pro-apoptotic and photo-protective properties (Pant *et al*., 2012). The first formal scientific reports on HS as therapeutic agents in modern Western clinical medicine are from 1957 in Hungary (Béres *et al*., 1957; Béres *et al*., 1958). Chinese medical literature also abounds with the therapeutic application of HS as those have consituted an important therapeutic class in Traditional Chinese Medicine since circa 3000 years. Humic substances have been since then clinically trialled and found to confer numerous beneficial features, e.g. they might protect the human body against blood coagulation or fibrinolysis as well as decreasing the effects of ionizing radiation (Klöcking and Björn, 2005).

Humic acids (HA) have good antioxidant activity and free radical scavenger ability (de Melo *et al*., 2016).

HS seem to have very few adverse effects (and even those only in specially hindered nutritional circumstances) and they are applicable as food and feed supplements (Murbach *et al*., 2020). Besides, HS have a wide agricultural and environmental use, e.g. as plant nutrients (Mosa *et al*., 2020). Nowadays humic matter is not only a human nutritional supplement, but it forms the basis of numerous feed additives in order to improve animal growth performance and health even by replacing antibiotic performance enhancers (Szabo *et al*., 2017). HS also have effects on gastrointestinal ulcer in pigs (Molnar, 2021) or rats (Li *et al*., 2011). HA forms a protective film on the surface of the ulcer in the stomach and helps cellular regeneration (Molnar, 2021). Fulvic acid (FA) combined with probiotics enhances the digestibility of phosphorous and metals. Moreover, it increased immune capability in a study conducted on a convincingly large number of pigs (Kunavue and Lien, 2012). HS as nutritional supplements have numerous beneficial effects on microelement and trace element homeostasis e.g. on iron and manganese homeostasis as proven in a rat model (Szabo *et al*., 2017) as well as copper and zinc homeostasis (Hullar *et al*., 2018). Some compounds of HS were found to have neuroprotective effects in animal studies, e.g., anthocyanin-containing gold-FA coated nanoparticles could prevent Alzheimer’s disease signs in rats (Kim *et al*., 2017). FAs not only reduce the assembly of tau proteins but also restore the original tau folding state (Cornejo *et al*., 2011). Some recent reports concluded that numerous compounds of HS, e.g. fulvic acid (Gnananath *et al*., 2020), are promising candidates for pharmaceutical use to enhance drug delivery of active pharmaceutical ingredients (APIs) (Jacob *et al*., 2019). HS have not only additional effects on health but they also have some therapeutic potential (Schepetkin *et al*., 2002). Fulvic acid has a clinical beneficial effect in chronic inflammatory diseases and diabetes (Winkler and Ghosh, 2018). The full HS fraction also has proven anti-inflammatory activity (Vucskits *et al*., 2010a; Vucskits *et al*., 2010b; Van Rensburg, 2015).

HS are known as antiviral agents for decades. Not long ago, a drink based on HS and plant extracts, branded as “Secomet V” was reported to possess broad spectral antiviral properties e.g. as anti-HIV, anti-poxvirus and anti-SARS activities (Kotwal *et al*., 2005). Numerous studies investigated and then reported antiviral features of HS e.g. against Coxsackie virus A9 (Klöcking and Sprössig, 1972), Herpes Simplex virus Type I. (Helbig *et al*., 1997), Influenza virus A/WSN/1933 (H1N1) (Lu *et al*., 2002), or HS being immune stimulators in Human Immunodeficiency virus-1 (HIV-1)-positive patients (Jooné *et al*., 2003), and tick-borne encephalitis virus infection (TBEV) (Orlov *et al*., 2019). A very recent high-quality study finds high antiviral activity of

HS compounds against HIV-1 (Zhernov *et al*., 2021). HS could also be regarded as pre-biotics inasmuch as they could provide beneficial effects in prophylaxis and therapy of coronaviral diseases (Tiwari *et al*., 2020). Given historical records and the scientific knowledge about the undisputedly high activity of HS against human viruses for more than fifty years, the authors attempted to investigate the possible antiviral activity of a commercially available nutritional supplement material. An *in vitro* quantitative real time Polymerase Chain Reaction (RT-qPCR)-based viral replication inhibition test was applied, using different dilutions of the investigated HS material.

In this material peat extract HS are enriched with Zn^2+^ ions, Se^-^ ions and ascorbic acid. Numerous studies confirmed that both ascorbic acid (Larenas-Linnemann *et al*., 2020; Grant *et al*., 2020), selenium ion (Bae and Kim, 2020; He *et al*., 2021) and zinc ions (Larenas-Linnemann *et al*., 2020; Quiles *et al*., 2020; te Velthuis *et al*., 2010), one by one, have anti-inflammatory effect and could reduce symptoms of COVID-19 infections. HS effectively bind metal ions as chelates (Saar and Weber, 1982; Kretzschmar and Christl, 2001; Bertoli *et al*., 2016), aromatic molecules and groups e.g. pyrene or Zn^2+^ ions, Se^-^ ions and ascorbic acid containg HS macromolecules could function with a nanoparticle-like colloidal behaviour (Jones and Bryan, 1998) which can enhance the antiviral activity of drug molecules (Chen and Liang, 2020) by virtue of their high surface to mass ratio.

## 2. Materials and Methods

The Test Article (TA) is a nutritional supplement base ingredient available commercially from its producer (ZnSeC-Humicin, Humic2000 Ltd, Budapest, Hungary) in powder or in solution and was commercially sourced by the authors. After sourcing, the purchased TA was sent to an independent commercial testing laboratory (Balint Analitika Ltd. Budapest) for the determination of ingredient concentrations using standardised, quality-assured and certified analytical test methods as prescribed in the Codex Alimentarius Hungaricus (Hungarian Food Book). All further in vitro cytopathogenicity and antiviral activity tests were performed at the Biosafety Level 3 (BSL-3) National Biosafety Laboratory (Budapest, Hungary).

### 2.1. Preparation of dilutions

The TA solution, a dark brown water-consistency fluid without any visible opalescence or sedimentation was prepared by the manufacturer. It was further centrifuged at 1300 rpm for 5 minutes to remove all eventual particulate inconsistencies. The TA solution originally contained 10g of ZnSeC-Humicin powder dissolved in 100 mL of distilled water. Next the supernatant was sterile-filtered through a 0.22 µm membrane filter. Then, a tenfold diluted stock was prepared from this filtrate using Dulbecco’s Modified Eagle Medium (DMEM, Gibco/ThermoFisher, CA, USA) cell culture medium which stock was also applied to study eventual cytopathogenic effects on a non infected VeroE6 cell culture. Further samples to be tested in the antiviral test were also diluted with DMEM. All sample dilutions were prepared from this filtered 1:10 diluted stock substance.

Final dilutions further tested thus were 100-fold, 500-fold, 1000-fold, 2000-fold, 5000-fold and 7000-fold.

### 2.2. Cell culture

VeroE6 cell line (ATCC® CRL-1586™) was maintained in DMEM cell culture medium with the addition of 10% m/m Fetal Bovine Serum (FBS, Gibco/ThermoFisher, CA, USA) supplemented with 50 U/ml penicillin and 5 µl/mL streptomycin. The studies after seeding the cells to plates, however, were further performed in FBS-free cell medium for vaccine production (VP-SFM, Gibco/ThermoFisher, CA, USA).

At first, cytopathogenicity of the stock solution *in vitro* was tested by adding 200 µl of TA stock solution to a plate’s each well with growing, uninfected VeroE6 cells and followed by incubation for 48 hours. For the *in vitro* effect tests per different TA dilutions, each TA dilution was incubated for 48 hours with the infected cells in the *in vitro* viral replication assay. 2×10^5^ cells/well VeroE6 cells were grown in 96-well culture plates and the listed various final dilutions of the TA were added to plate wells (cf. 2.1).

### 2.3. In vitro viral replication assay

SARS-CoV-2 virus B1.1.7 Variant of Concern (“Alpha VOC”), stocked in frozen form at -80°C at the National Biosafety Laboratory isolated from Hungarian samples in January 2021 was cultured in VeroE6 cells and divided into aliquots, which were used for the assays.

Prior to this, the amounts of plaque forming unit (PFU) and Tissue Culture Infective Dose at 50% (TCID_50_) were determined and the corresponding amounts were set aside as mentioned.

### 2.4. Test for the inhibition of SARS-CoV-2 replication

To detect viral replication, we examined the supernatants from VeroE6 cells cultured for 48 h in the presence of the TA dilutions. The assay was performed using qRT-PCR and the QuantiNova Probe RT-PCR kit (Qiagen, Düsseldorf, Germany), and the LightCycler 480 Real-Time PCR System (Roche, Basel, Switzerland). RNA extraction was performed using the Roche High Pure Viral RNA Kit (Roche, Basel, Switzerland) according to the manufacturer’s instructions. An additional washing step with a PerkinElmer Chemagic (PerkinElmer, USA) automated extraction instrument was also included (with wash buffer) to remove potential inhibitors of PCR.

We then targeted the N-(nucleoprotein) gene of SARS-CoV-2 virus with Roche Lightcycler 480 RNA Master Hydrolysis Probes (Roche Basel, Switzerland) with primers developed in house, using the Lightcycler 2.0 (Roche) platform. Primers for RNA virus detection were previously validated by the Biosafety Laboratory (troughout at BSL-3) through and in collaboration with, the European Research Infrastructure on Highly Pathogenic Agents (ERINHA).

In the assay VeroE6 cells were incubated using 96-well plate until a monolayer was formed. Each of the TA dilutions to be tested was suspended (vortexed for 30 seconds) and then pipetted immediately. The same 200 µl volume of the TA (throughout all the dilutions) was added to the solution containing the volume of SARS-CoV-2 virus corresponding to 100 TCID_50_, to detect their virus inhibiting/antiviral ability (see below).

For the citotoxicity assay a spectrophotometer (Omni cell adhesion light spectrometer, Cytosmart, Netherlands) was used. The percentage rate of surface covered by monolayers and of transparent surface (plaques appearing in the absence of cells killed by the virus) in all of the wells of the 96-well cell culture platform were determined.

All tests were made in triplicates. From these the relative parameters of viral inhibition were calculated compared to the viral effect in an infected, untreated cell culture free of TA.

#### 2.4.1. Test protocol

Different experimental groups were applied: the “virus control” group was a positive control, untreated with any TA, to determine the effects of the virus infection. The inhibitory effect of TA dilutions was compared to the result of this group.

The cell control group was a negative control, where the effect of just the culturing activities on virus-free cells was measured; the group was also free of the TAs.

Dilution test groups were set up with each TA dilution added to the wells and incubated with the virus as follows. First, the active ingredients were mixed in a 1:1 volumetric ratio with 100 TCID_50_ SARS-CoV-2 virus suspension. The second, infection phase applied a SARS-CoV-2 virus suspension of 100 TCID_50_ (0.1 multiplicity of infection, MOI), mixed with the dilutions of the TA in a volumetric ratio of 1:1, which mixture was put on the cell culture and then incubated at 37 °C in 5% CO_2_ atmosphere for 2 hours in the absence of FBS.

After the infection phase, the infectious virus-containing mixture was removed with an automatic pipette. Immediately after this, an 1:1 mixture of VP-SFM medium and the appropriate DMEM medium dilution of the TA was added to the cell cultures in equally 200 microliter volumes. Incubation with this active ingredient-treated medium lasted for 48 hours.

After 48 hours, the virus-containing medium containing the TA was removed from the cells. The removed viral supernatant was ultracentrifuged at 16.000 g and the supernatant was further measured for viral RNA content: The supernatant was subjected to PCR assay to determine the viral RNA content, as previously mentioned, for the N-gene. The supernatant RNA contents were quantified for gene copy numbers and compared to the number of copies measured in the untreated, not inhibited SARS-CoV-2 infected cell cultures to calculate the inhibitory effect.

#### 2.4.2 Mathematical analysis of results

Viral N-gene copy numbers obtained in the triplicates both for inhibitory TA and untreated virus control wells were used as input data. The aim of mathematical data analysis was to determine the inhibitory dilution level at which the number of viral N-gene copy numbers declines to the 50% of virus control (50% inhibitory dilution, Dilution_50_). Then, this dilution value would serve as the basis to calculate the 50% inhibitory “dose”: the molar concentration for each ingredient (ID_50_)

The copy numbers were the input for a sigmoidal curve fit using our own in-house developed algorithm and software in Octave mathematical environment.

The sigmoidal function was generalised as **(Eq. 1)**

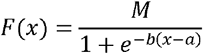

where

**M** is the maximal viral copy number (at infinite dilution of the added TA i.e. with only infinitesimally small TA content of the plate wells)

**a** is the dilution value of the TA at 50% of M (Dilution_50_)

**b** is the derivate of intermediate transmission.

We then characterised the curve fit and calculated the value of parameter **a** and its 95% confidence interval. As a further step towards obtaining ID_50_ values, the active ingredient molar concentrations were determined from the obtained Dilution_50_ value using the known composition, the applied volume per well, and molar masses of ingredients. This allows the comparison of the TA inhibitory effect, to other in vitro inhibition results of different other molecules or extracts.

## 3. Results and discussion

In our viral replication inhibition asay, two main findings are reported. The SARS-CoV-2 virus B1.1.7 VOC was used as infective agent (chapter 2.3).

1. Neither qualitative nor quantitative cytotoxicity was observed in any of the tested TA dilutions on VeroE6 cells.
2. A dose-dependent inhibition was demonstrated.

The active components of the commercially sourced “ZnSeC-Humicin” nutritional supplement material used for the experiment and their concentrations are summarized in Table 1. The original HS components (65.9 m/m% FA and 0.5 m/m% HA) are compounded with high concentration of Vitamin C (about 40 g/100g) and lower concentration of metal ions (60900 mg/kg Zinc-ion and 23.2 mg/kg Selenium ion according to their recommended daily intake values in human.

**Table 1.**
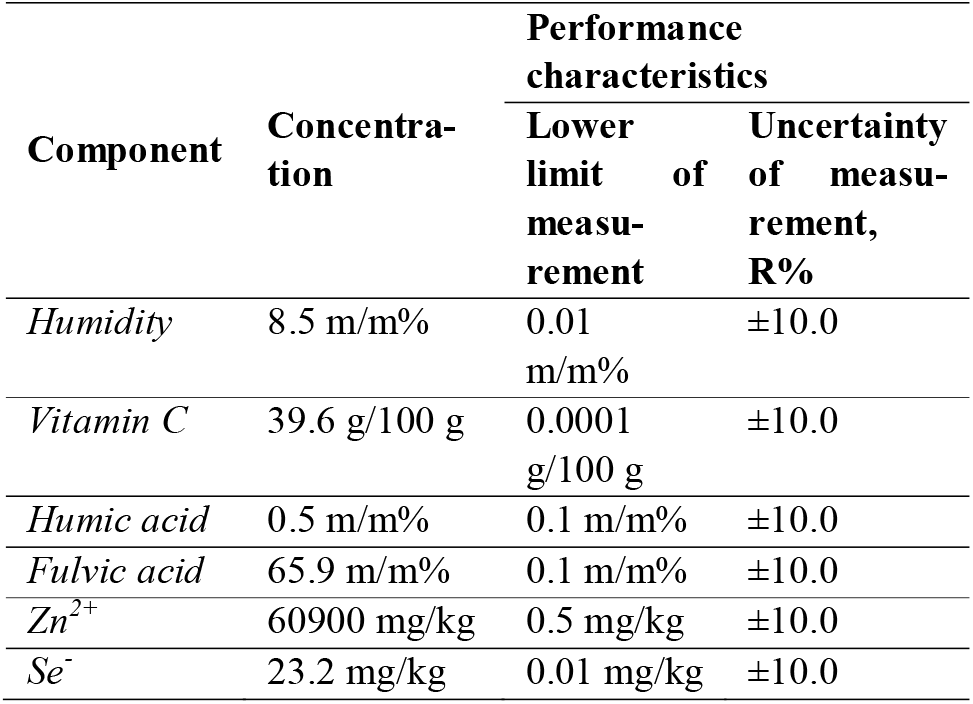
Ingredients of the tested “ZnSeC-Humicin” powder nutritional supplement material.

The obtained dilution value and other inhibition curve characteristics are presented in ***Table 2***.

**Table 2.** Characteristics of the inhibition curve of the Test Article against SARS-CoV-2 Alpha VOC

| Parameter | Fit Value | Confidence interval at 95% |
| --- | --- | --- |
| **M** | $7.23 \times 10^4$ | $(51.52 \times 10^4, 94.95 \times 10^4)$ |
| **a** | **1739.98** | $(621.80, 2858.00)$ |
| **b** | $4.84 \times 10^{-3}$ | $(-14.12 \times 10^{-3}, 23.80 \times 10^{-3})$ |

Test results show that the humic substances compounded with Zn-Se-ascorbic acid (ZnSeC-Humicin) seemingly exert a significant inhibitory effect on SARS-COV-2 virus replication (Figure 2.) A sigmoidal curve (red continuous line) could be fitted to the triplicate copy number data points (blue squares) for determination of the mean inhibitory dose (ID_50_) value of the tested HS material. Figure 2. shows a semi logarithmic scale transformation of the sigmoidal function. It was found that the Dilution_50_ value of ZnSeC-Humicin stock solution is close to 2000 (1740) times dilution.

**Figure 2.**
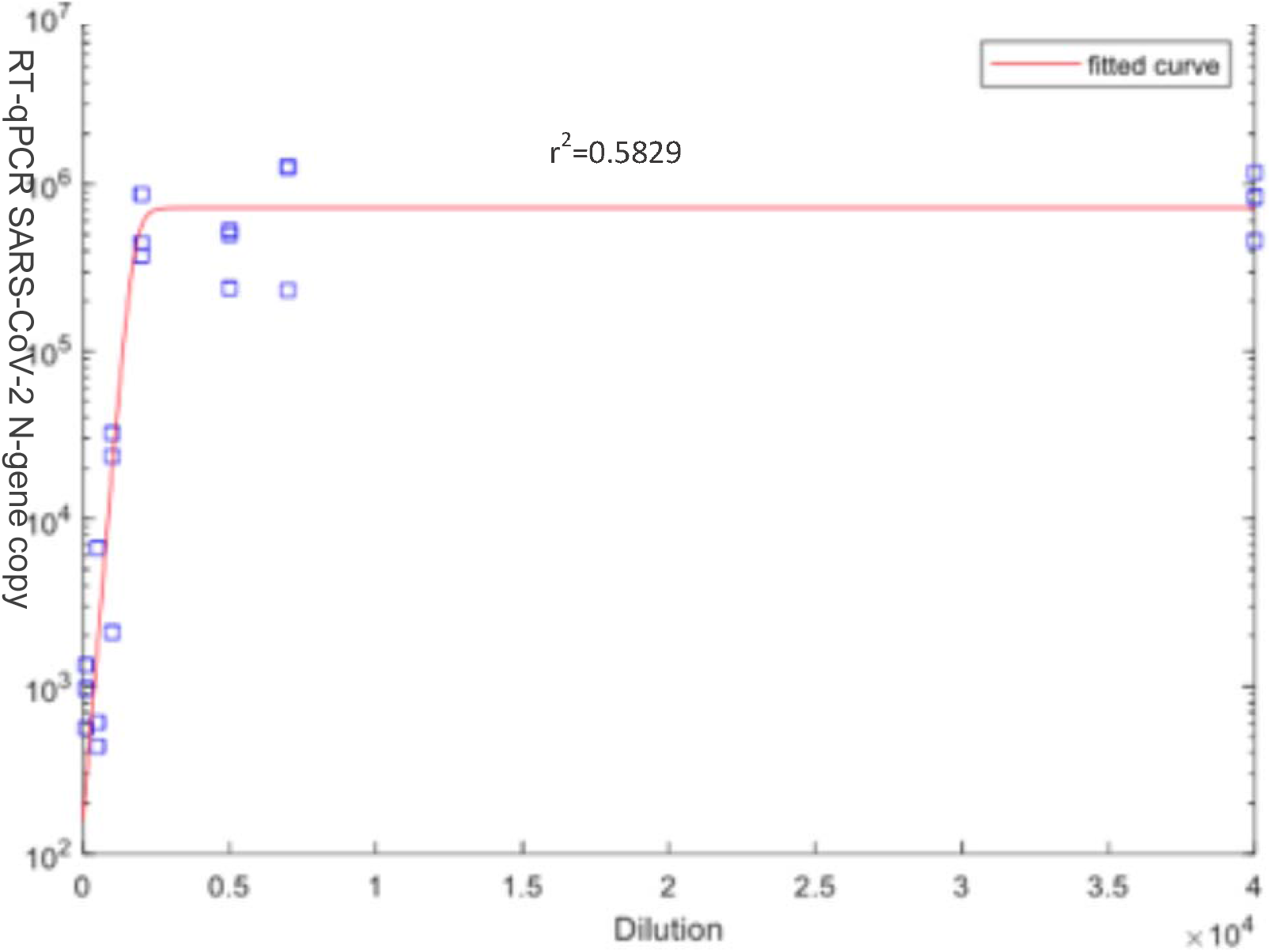
Inhibitory effect of HS compared with Zn-Se-ascorbic acid complex (ZnSeCHS) for SARS-CoV-2 virus B1.1.7 variant of concern (“Alpha VOC”) in Vero F6 cell culture.

Molar concentrations of ingredients of ZnSeC-Humicin at dilution in the ID_50_ value are usually in a 10^−9^ M (pM) concentration range. The measured relative humidity of the powder was 8.5% m/m thus data were corrected to dry weight in the table. A very low, 70.80 picomol/L concentration of FA and a similarly also very low, femtomolar (49.00-409.00 fM) concentration of HA with 610.00 pM ascorbic acid, 251.00 pM Zn^2+^ and 7.28 pM of Se^-^ can reduce to half the copy numbers of SARS-CoV-2 virus B1.1.7 variant of concern (“Alpha VOC”) (Table 3.)

**Table 3.**
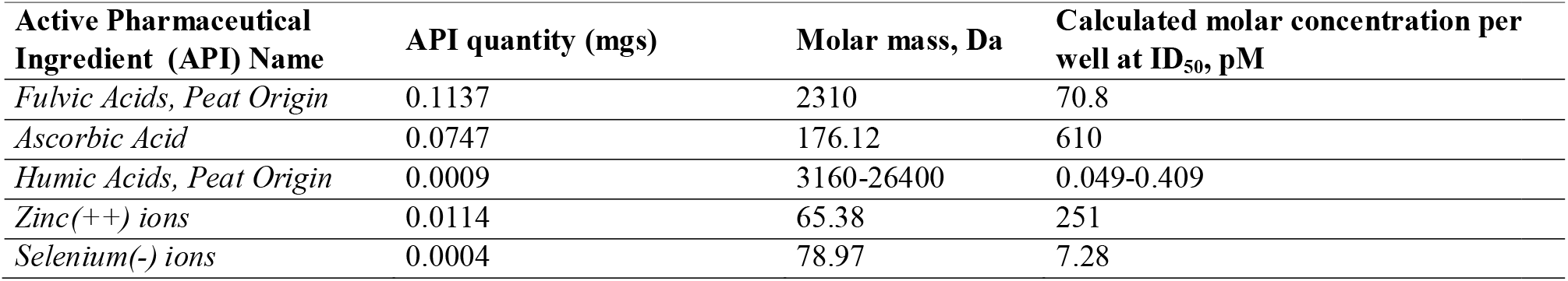
Calculated API quantities for the observed ID_50_ effect 1740-fold dilution, measured in the applied 200 microliter volumes.

## 4. Conclusions

This proof of concept study shows that the Test Article, a peat extract HS and Zn/Se based nutritional supplement possesses measurable, dose-dependent and robust in vitro inhibitory effects on SARS-CoV-2 infection and replication.

Normalized to API molar masses of Humic Acid, a hundred femtomolar range is apparent, while for Selenium, a picomolar range, for Fulvic Acids, the inhibition constants are in the decimal picomolar range while for Ascorbic Acid and Zinc in the hundred picomolar range.

Both fulvic acids and Zinc ions have been proven to exert powerful and dose dependent antiviral effects against HIV, influenza- and coronaviruses. By potentiating the effects of FA with an ascorbic acid complex that can also bind Zn ions, and by increasing cellular antioxidant protection turnover using Selenium for glutathione peroxidase, we postulate a synergistic effect of these ingredients. The main antiviral / replication inhibitory effects are probably exerted by FA and Zn^++^ ions. As there are now multiple reports on intestinal and even stool-based replication of SARS-CoV-2, the hypothesis of this proof-of-concept study should be further tested in pre-clinical animal models of peroral administration, and in clinical trials. These in vitro results are by no means a direct sign of human clinical applicability with a therapeutic intent, but they do direct us for further investigations on the effects of materials possessing proven in vitro antiviral activity used in combination.

The picomolar concentration ranges also indicate that further *in vivo* studies should be conducted. These values are at least one magnitude lower than the *in vitro* inhibition values of almost all other available antiviral molecules.

## Acknowledgements

This work was supported by the Higher Education Institutional Excellence Program of the Ministry of Human Resources in Hungary, within the framework of the Therapeutic Development thematic program of Semmelweis University, the European Union’s Horizon 2020 research and innovation program under grant agreement No 739593: HCEMM, supported by EU Program: H2020-EU.4.a. This work was also partly funded by grants from the Hungarian National Research, Development and Innovation Office (Thematic Excellence Program, TKP-BIOImaging, financed under the 2020-4.1.1-TKP2020 funding scheme and Investment to the Future 2020.1.16-Jövő-2021-00013).

## Notes

### Competing Interest Statement

The authors have declared no competing interest.

### Summary of Updates

The previously published (misordered) author list was corrected.

## References

Alvarez-Puebla A R n, Garrido J J and Aroca R F 2004 Surface-Enhanced Vibrational Microspectroscopy of Fulvic Acid Micelles Analytical Chemistry 76 7118–25

Alvarez-Puebla R A, Valenzuela-Calahorro C and Garrido J J 2006 Theoretical study on fulvic acid structure, conformation and aggregation. A molecular modelling approach Sci Total Environ 358 243–54

Asakawa D, Kiyota T, Yanagi Y and Fujitake N 2008 Optimization of Conditions for High-Performance Size-Exclusion Chromatography of Different Soil Humic Acids Analytical Sciences 24 607–13

Bae M and Kim H 2020 The role of vitamin C, vitamin D, and selenium in immune system against COVID-19 Molecules 25 5346

Béres T, Kabdebó S, Ferenc K, Nemeséri L, Szélsy A and Visy L 1957 [Studies on therapeutic application of fulvic acids, with special regard to their liver protecting function. In Hungarian.] Magy Állatorv Lapja 12 351–2

Béres T, Király I, Bóna E, Lővei E and Róbert S 1958 Tőzeg-fulvósavval szerzett therapiás tapasztalataink Orvosi Hetilap 17 567

Bertoli A C, Garcia J S, Trevisan M G, Ramalho T C and Freitas M P 2016 Interactions fulvate-metal (Zn(2)(+), Cu(2)(+) and Fe(2)(+)): theoretical investigation of thermodynamic, structural and spectroscopic properties Biometals 29 275–85

Chen L and Liang J 2020 An overview of functional nanoparticles as novel emerging antiviral therapeutic agents Mater Sci Eng C Mater Biol Appl 112 110924

Chin Y-P, Aiken G and O’Loughlin E 1994 Molecular weight, polydispersity, and spectroscopic properties of aquatic humic substances Environmental science & technology 28 1853–8

Chon K, Cho J and Shon H K 2013 Advanced characterization of algogenic organic matter, bacterial organic matter, humic acids and fulvic acids Water Sci Technol 67 2228–35

Cornejo A, Jiménez J M, Caballero L, Melo F and Maccioni R B 2011 Fulvic acid inhibits aggregation and promotes disassembly of tau fibrils associated with Alzheimer’s disease Journal of Alzheimer’s disease 27 143–53

de Melo B A, Motta F L and Santana M H 2016 Humic acids: Structural properties and multiple functionalities for novel technological developments Mater Sci Eng C Mater Biol Appl 62 967–74

Ghosal S, Baumik S and Chattopadhyay S 1995 Shilajit induced morphometric and functional changes in mouse peritoneal macrophages Phytotherapy Research 9 194–8

Ghosal S, Lal J, Singh S K, Goel R K, Jaiswal A K and Bhattacharya S K 1991 The need for formulation of Shilajit by its isolated active constituents Phytotherapy research 5 211–6

Gnananath K, Nataraj K S, Rao B G, Kumar K P, Mahnashi M H, Anwer M K, Umar A, Iqbal Z and Mirza M A 2020 Exploration of fulvic acid as a functional excipient in line with the regulatory requirement Environ Res 187 109642

Grant W B, Lahore H, McDonnell S L, Baggerly C A, French C B, Aliano J L and Bhattoa H P 2020 Evidence that Vitamin D Supplementation Could Reduce Risk of Influenza and COVID-19 Infections and Deaths Nutrients 12

He L, Zhao J, Wang L, Liu Q, Fan Y, Li B, Yu Y-L, Chen C and Li Y-F 2021 Using nano-selenium to combat Coronavirus Disease 2019 (COVID-19)? Nano Today 36 101037

Helbig B, Klockinq R and Wutzler P 1997 Anti-herpes simplex virus type 1 activity of humic acid-like polymers and their o-diphenolic starting compounds Antiviral Chemistry & Chemotherapy 8 265–73

Hullar I, Vucskits A V, Berta E, Andrasofszky E, Bersenyi A and Szabo J 2018 Effect of fulvic and humic acids on copper and zinc homeostasis in rats Acta Vet Hung 66 40–51

Jacob K K, Kj P P and N C 2019 Humic Substances as a Potent Biomaterials for Therapeutic and Drug Delivery System-a Review International Journal of Applied Pharmaceutics 1–4

Jones M N and Bryan N D 1998 Colloidal properties of humic substances Advances in Colloid and Interface Science 78 1–48

Jooné G K, Dekker J and Jansen Van Rensburg C E 2003 Investigation of the Immunostimulatory Properties of Oxihumate Zeitschrift für Naturforschung C 58 263–7

Kim M J, Rehman S U, Amin F U and Kim M O 2017 Enhanced neuroprotection of anthocyanin-loaded PEG-gold nanoparticles against Abeta1-42-induced neuroinflammation and neurodegeneration via the NF-KB/JNK/GSK3beta signaling pathway Nanomedicine 13 2533–44

Klöcking R and Björn H 2005 Biopolymers for Medical and Pharmaceutical Applications, (Weinheim WILEY-VCH Verlag GmbH & C. KGaA.) pp 3–16.

Klöcking R and Sprössig M 1972 Antiviral properties of humic acids Experientia 28 607–8

Kotwal G J, Kaczmarek J N, Leivers S, Ghebremariam Y T, Kulkarni A P, Bauer G, De Beer C, Preiser W and Mohamed A R 2005 Anti-HIV, anti-poxvirus, and anti-SARS activity of a nontoxic, acidic plant extract from the Trifollium species Secomet-V/anti-vac suggests that it contains a novel broad-spectrum antiviral Ann N Y Acad Sci 1056 293–302

Kretzschmar R and Christl I 2001 Proton and metal cation binding to humic substances in relation to chemical composition and molecular size SPECIAL PUBLICATION-ROYAL SOCIETY OF CHEMISTRY 273 153–64

Kumar Gautam R, Navaratna D, Muthukumaran S, Singh A, Islamuddin and More N 2021: IntechOpen) Kunavue N and Lien T F 2012 Effects of Fulvic Acid and Probiotic on Growth Prerformance, Nutrient Digestibility, Blood Parameters and immunity of Pigs.pdf> J. Anim.Sci. Adv. 2 711–21

Lag J, Hadas A, Fairbridge R W, Muñoz J C N, Pombal X P, Cortizas A M and Almendros G 2008 Encyclopedia of Soil Science, ed W Chesworth (Dordrecht: Springer Netherlands) pp 315–23

Larenas-Linnemann D, Rodriguez-Perez N, Arias-Cruz A, Blandon-Vijil M V, Del Rio-Navarro B E, Estrada-Cardona A, Gereda J E, Luna-Pech J A, Navarrete-Rodriguez E M, Onuma-Takane E, Pozo-Beltran C F and Rojo-Gutierrez M I 2020 Enhancing innate immunity against virus in times of COVID-19: Trying to untangle facts from fictions World Allergy Organ J 13 100476

Li Y-M, Li B-C, Li P, Liu J-Z, Cui J-L and Mei Z-Q 2011 Effects of Na-FA on gastrointestinal movement and gastric ulcer in mice Zhong yao cai= Zhongyaocai= Journal of Chinese medicinal materials 34 1565–9

Lieke T, Steinberg C E W, Pan B, Perminova I V, Meinelt T, Knopf K and Kloas W 2021 Phenol-rich fulvic acid as a water additive enhances growth, reduces stress, and stimulates the immune system of fish in aquaculture Scientific Reports 11

Lu F J, Tseng S N, Li M L and Shih S R 2002 In vitro anti-influenza virus activity of synthetic humate analogues derived from protocatechuic acid Archives of Virology 147 273–84

Mishra T, Dhaliwal H S, Singh K and Singh N 2019 Shilajit (Mumie): Current Status of Biochemical, Therapeutic and Clinical Advances Current Nutrition & Food Science 15 104–20

Molnar D 2021 The beneficial effects of humic acid on gastric ulcers in pigs International Pig Topics 34 9

Mosa A, Taha A and Elsaeid M 2020 Agro-environmental applications of humic substances: A critical review Egyptian Journal of Soil Science 0 0-

Murbach T S, Glavits R, Endres J R, Clewell A E, Hirka G, Vertesi A, Beres E and Pasics Szakonyine I 2020 A toxicological evaluation of a fulvic and humic acids preparation Toxicol Rep 7 1242–54

Nebbioso A and Piccolo A 2013 Molecular characterization of dissolved organic matter (DOM): a critical review Analytical and Bioanalytical Chemistry 405 109–24

Orlov A A, Zherebker A, Eletskaya A A, Chernikov V S, Kozlovskaya L I, Zhernov Y V, Kostyukevich Y, Palyulin V A, Nikolaev E N, Osolodkin D I and Perminova I V 2019 Examination of molecular space and feasible structures of bioactive components of humic substances by FTICR MS data mining in ChEMBL database Scientific Reports 9

Pant K, Singh B and Thakur N 2012 Shilajit: A Humic Matter Panacea for Cancer International Journal of Toxicological and Pharmacological Research 4 17–25

Quiles J L, Rivas-Garcia L, Varela-Lopez A, Llopis J, Battino M and Sanchez-Gonzalez C 2020 Do nutrients and other bioactive molecules from foods have anything to say in the treatment against COVID-19? Environ Res 191 110053

Saar R A and Weber J H 1982 Fulvic acid: modifier of metal-ion chemistry Environmental science & technology 16 510A–7A

Schellekens J, Buurman P, Kalbitz K, Zomeren A V, Vidal-Torrado P, Cerli C and Comans R N J 2017 Molecular Features of Humic Acids and Fulvic Acids from Contrasting Environments Environmental Science & Technology 51 1330–9

Schepetkin I, Khlebnikov A and Kwon B S 2002 Medical drugs from humus matter: Focus on mumie Drug Development Research 57 140–59

Stevenson F J 1994 Humus chemistry: genesis, composition, reactions: John Wiley & Sons)

Sutton R and Sposito G 2005 Molecular structure in soil humic substances: the new view Environmental science & technology 39 9009–15

Szabo J, Vucskits A V, Berta E, Andrasofszky E, Bersenyi A and Hullar I 2017 Effect of fulvic and humic acids on iron and manganese homeostasis in rats Acta Vet Hung 65 66–80

te Velthuis A J, van den Worm S H, Sims A C, Baric R S, Snijder E J and van Hemert M J 2010 Zn(2+) inhibits coronavirus and arterivirus RNA polymerase activity in vitro and zinc ionophores block the replication of these viruses in cell culture PLoS Pathog 6 e1001176

Tiwari S K, Dicks L M T, Popov I V, Karaseva A, Ermakov A M, Suvorov A, Tagg J R, Weeks R and Chikindas M L 2020 Probiotics at War Against Viruses: What Is Missing From the Picture? Front Microbiol 11 1877

Van Rensburg C E J 2015 The Antiinflammatory Properties of Humic Substances: A Mini Review Phytotherapy Research 29 791–5

Vucskits A, Hullár I, Bersenyi A, Andrásofszky E, Tuboly T and Szabó J 2010a Effect of fulvic acid and humic acid. Part 1. Economic indexes, immunostimulant effect Magyar Állatorvosok Lapja 132 278–84

Vucskits A V, Hullár I, Bersényi A, Andrásofszky E, Kulcsár M and Szabó J 2010b Effect of fulvic and humic acids on performance, immune response and thyroid function in rats Journal of Animal Physiology and Animal Nutrition 94 721–8

Winkler J and Ghosh S 2018 Therapeutic Potential of Fulvic Acid in Chronic Inflammatory Diseases and Diabetes J Diabetes Res 2018 5391014

Zhernov Y V, Konstantinov A I, Zherebker A, Nikolaev E, Orlov A, Savinykh M I, Kornilaeva G V, Karamov E V and Perminova I V 2021 Antiviral activity of natural humic substances and shilajit materials against HIV-1: Relation to structure Environ Res 193 110312

